# Hand and face somatotopy shown using MRI-safe vibrotactile stimulation with a novel Soft Pneumatic Actuator (SPA)-Skin interface

**DOI:** 10.1101/2021.10.27.465604

**Authors:** Sanne Kikkert, Harshal A. Sonar, Patrick Freund, Jamie Paik, Nicole Wenderoth

## Abstract

The exact somatotopy of the human facial representation in the primary somatosensory cortex (S1) remains debated. One reason that progress has been hampered is due to the methodological challenge of how to apply automated vibrotactile stimuli to face areas in a manner that is: 1) reliable despite differences in the curvatures of face locations; and 2) MR-compatible and free of MR-interference artefacts when applied in the MR head-coil. Here we overcome this challenge by using soft pneumatic actuator (SPA) technology. SPAs are made of a soft silicon material and can be in- or deflated by means of airflow, have a small diameter, and are flexible in structure, enabling good skin contact even on curved body surfaces (as on the face). To validate our approach, we first mapped the well-characterised S1 finger layout using this novel device and confirmed that tactile stimulation of the fingers elicited characteristic somatotopic finger activations in S1. We then used the device to automatically and systematically deliver somatosensory stimulation to different face locations. We found that the forehead representation was least distant from the representation of the hand. Within the face representation, we found that the lip representation is most distant from the forehead representation, with the chin represented in between. Together, our results demonstrate that this novel MR compatible device produces robust and clear somatotopic representational patterns using vibrotactile stimulation through SPA-technology.

## Introduction

Over the past decades, somatotopic mapping provided us with an increasingly better understanding of how the body is represented in the brain. However, providing tactile stimulation in an MR environment remains challenging. Experimenters commonly use an active movement paradigm (Kolasinski et al., 2016; Root et al., 2021; Schellekens et al., 2018; Zeharia et al., 2015) or manual stroking of body parts (Martuzzi et al., 2014; Sanders et al., 2019; Van Der Zwaag et al., 2015) to probe somatotopic representations. However, both paradigms are limited: movement execution does not allow to study the representation of tactile stimuli in isolation and distinguishing between body parts that are not able to move independently (e.g., certain parts of the face or toes) is impossible. Manual stroking induces experimenter dependent spatiotemporal variance since the intensity, timing, and body part coverage of manual stimulations may not be consistent and precise. Furthermore, there are practical challenges when multiple body parts need to be stroked simultaneously, or if space is limited (e.g. when using a narrow head- or body coil). The usage of mechanical vibrotactile devices provides an opportunity to overcome these limitations. However, common vibrotactile elements contain metals or electrical circuits that are mostly not compatible with the MR environment (Yu and Riener, 2006). Stimulating body parts in (or close to) the MR head coil is especially technically demanding given the narrow geometry of the head coil and the safety constraints and imaging artefacts induced by metal components even if they are small.

Commercially available piezoelectric and piezoceramic devices are able to deliver vibrotactile stimulation at high-frequency ranges at a fixed amplitude. However, these devices mostly still contain some metal components and lead electrical wires inside the scanner (Puckett et al., 2017; Sanchez-Panchuelo et al., 2010). While active RF shielding is sufficient to prevent heating by RF pulses when such stimulators are placed far from the head coil such as on the fingers, they could still induce unwanted signal interference when placed closer to, or inside, the MR head coil. Pneumatically driven or air puff devices circumvent these issues and can be made of non-metallic MR compatible materials.

The pneumatic devices that have been developed for safe delivery of tactile stimulation to multiple stimulation sites inside the MR head coil are however limited. The Dodecapus was amongst the first pneumatically driven devices build to apply automated tactile stimulation via air puffs to a range of face locations (Huang and Sereno, 2007). The device was extended to deliver punctate tactile stimuli to the face using Von Frey filaments with a high spatiotemporal accuracy (Dresel et al., 2008). However, in these setups stimulation could only be applied to parts of the face that are exposed (i.e., that are not covered by the head coil). Later, flexible plastic tubes were attached to a facial mask (custom moulded for each subject) to deliver air puffs to locations of the face that are covered by the head coil, allowing for a more complete picture of face somatotopy (Chen et al., 2017; Huang et al., 2017). However, this face mask may not fit in head coils with more narrow geometry and when using participants with larger head sizes. Furthermore, providing air puffs close to the mouth, nose, or eyes may not be comfortable for participants. More recently, the GALILEO Somatosensory^TM^ was developed. This device can provide tactile stimulation through pressure dynamics in individual stimulators with high spatiotemporal control (Custead et al., 2017). While this device is very promising and allows the study of stimulation velocity, the stimulators are 6mm in height and may not fit in head coils with very narrow geometry. Furthermore, it may not be easy to attach the stimulators to highly curved body surfaces and the device is rather expensive.

An optimal vibrotactile device for MRI usage comprehensively consists of MR-safe materials, is safe to place on the skin, and does not induce any MR interference artefacts even inside the MR head coil. Furthermore, it should allow a controllability over vibration frequency and amplitude, enabling both sub- and suprathreshold vibrotactile stimulations. The stimulators themselves should be flexible in nature, such that they can be placed on skin surfaces with different curvatures, and small and narrow to provide focal stimulation in narrow head coils that are typical for ultra-high field imaging. Lastly, the device should be portable and cost-efficient.

Here we present a newly developed tactile stimulation device that delivers all the characteristics of the aforementioned “optimal vibrotactile” device. This novel platform provides focal, suprathreshold vibrotactile stimulation to the face (even inside a narrow head coil) or to other parts of the body inside an MRI scanner. The soft pneumatic actuator (SPA)- skin does not contain any metals at the contact point and still provides thorough control over actuation frequencies (Sonar et al., 2019; Sonar and Paik, 2016). SPAs are made of a soft MR-safe silicone (Dragon Skin 30®, Smooth On Inc., USA) and can be inflated or deflated by varying internal pressure. SPA-skins are extremely versatile in their geometric dimensions and material choices: the presented prototype measures under 1 mm thin, with a 1.4 cm diameter actuation point. It is soft and flexible in structure, enabling compliant skin contact even on curved body surfaces (as on the face). Furthermore, the low profile design of the SPA-skin allows for usage in the narrow geometry of the MR head coil and in narrow bores that are typical for ultra-high field MRI environments. The choice of materials with matching mechanical compliance with human skin improves the optimal transfer of tactile feedback while maintaining mechanical transparency when worn.

We demonstrated the feasibility of using the SPA-skin setup for somatotopic mapping in an fMRI study. By doing so we not only provide a methodological but also a scientific advance as our device allowed us to detail the somatotopic layout of the face via an automated face stimulation paradigm. While the gross representation of body parts and fingers in S1 is largely agreeable across studies, results on the exact somatotopic mapping of the face have been mixed. Indeed, contrasting work reports both an inverted (Servos et al., 1999; Yang et al., 1993) and upright somatotopic face representation (Huang and Sereno, 2007; Penfield and Rasmussen, 1950; Root et al., 2021; Roux et al., 2018; Sato et al., 2005; Schwartz et al., 2004). Yet other studies reported more mixed face representations following an ‘onion skin model’ where the nose was represented more inferior in S1 than stimulations on the lower jaw or above the eye (DaSilva et al., 2002; Moulton et al., 2009). To detail the full somatotopic layout of the face an automated face stimulation paradigm and both univariate analysis and multivariate analysis methods that are sensitive to representational overlaps (i.e., representational similarity analysis) are needed.

We hypothesized that tactile SPA stimulation would yield robust localised activations in somatosensory areas. We further hypothesized that tactile finger stimulation would elicit characteristic somatotopic finger activations in the primary somatosensory cortex. Lastly, we investigated the somatotopic layout of the face, using the device to automatically and systematically deliver somatosensory stimulation inside the head coil.

## Material and methods

### Participants

Seventeen healthy participants (mean age ± s.e.m. = 37.1 ± 4.3; 7 females; 2 left-handers) participated in this study. Ethical approval was granted by the Kantonale Ethikkommission Zürich (EK-2018-00937) and written informed consent was obtained prior to study onset. This study is registered on clinicaltrials.gov under the number NCT03772548. As our fingers stimulation experiment was meant as a mere validation of the device and was expected to reveal convincing data in a limited number of participants, we only tested 8 participants for this feasibility aspect of the study. Face somatotopy was tested in all participants.

### SPA-skin stimulator design

We developed a novel soft skin like interface (Figure 1): i.e., a SPA-skin that can be driven using modulated pneumatic pressure and is capable of generating a plethora of actuation < 100 Hz and can be complemented by integrated sensors (Sonar et al., 2019; Sonar et al., 2019; Sonar and Paik, 2016). The SPA-skin is fabricated using silicone (Dragon Skin 30, Smooth-On Inc., USA) that has similar stiffness as the human skin and provides optimal comfort, as is necessary for long-duration fMRI studies (Figure 2). The actuator layer, i.e., the SPA, consists of an elastomeric membrane that can be pneumatically inflated with a pressure input (Sonar and Paik, 2016; see Figure 2B). This actuator is fabricated with three thin layers with a total thickness of < 1mm: a middle flexible mask layer (50μm) to define the actuator’s shape that is sandwiched between two silicone layers. The masking layer is laser machined to obtain the desired shape and is then laminated onto the bottom silicone layer to be encapsulated by a thin top silicone layer. The polypropylene mask adheres to the bottom silicone layer and ensures that, upon inflation, the top membrane deforms. It is important to ensure a proper grounding with the skin and attaching the SPA-skin properly at a given location without obstructing the inlet pneumatic flow lines.

**Figure 1:**
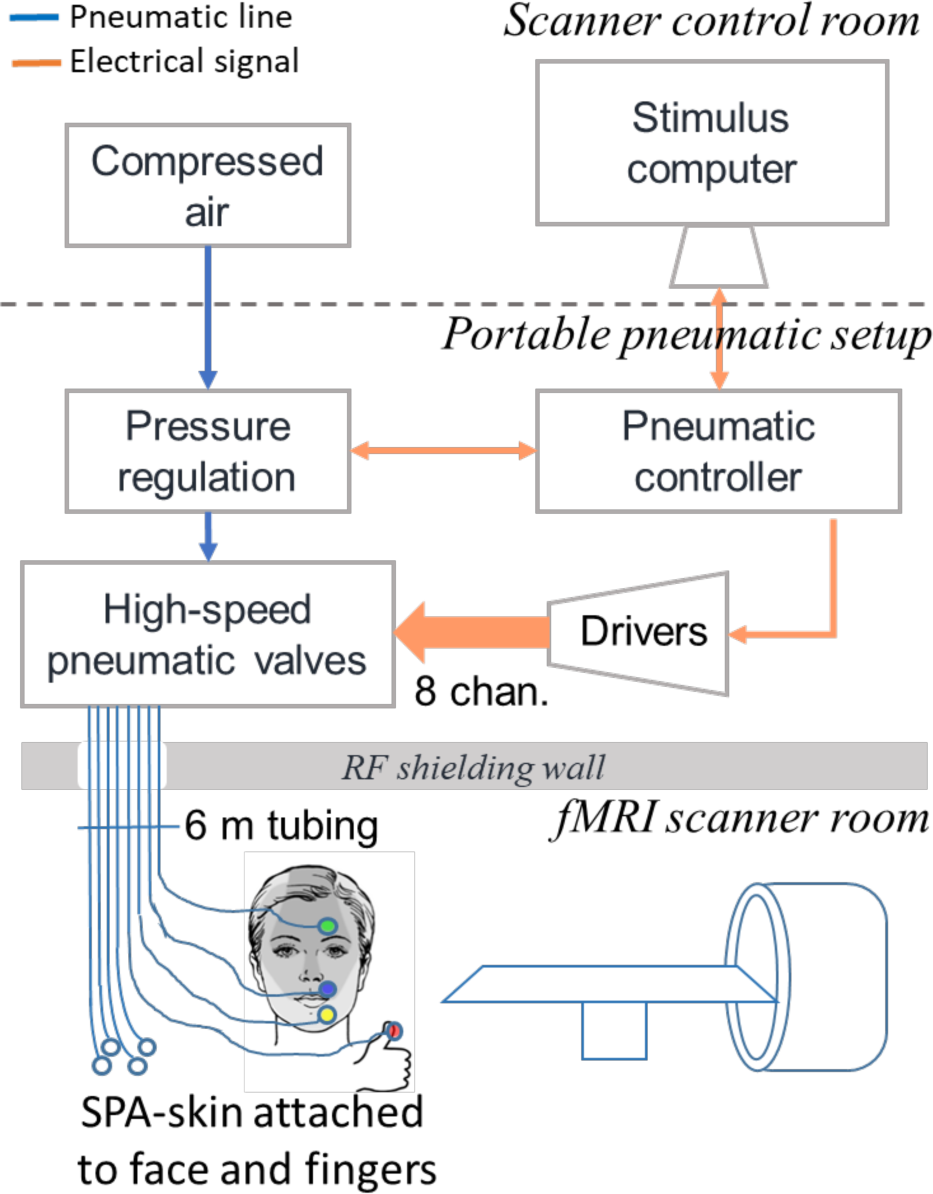
Schematic illustration of the SPA setup. The pneumatic controller was composed of the following components: a stimulus computer, a portable air compressor, a portable pneumatic setup, and pneumatic tubes. Stimulation intensity and frequency could be controlled using a stimulus computer in the scanner control room. An air compressor placed in the scanner control room provided airflow to a pressure regulator that was controlled by the microcontroller in the portable pneumatic setup. The stimulation intensity could be adjusted using the pressure regulator and an array of high-speed pneumatic valves which were driven through high-power switches and inputs from the micro-controller. A 50mm diameter hole in the RF shielding wall then carried the 5-6m long and 4mm thick tubes to the SPAs that were placed on the face or the fingers in the fMRI scanner room.

**Figure 2:**
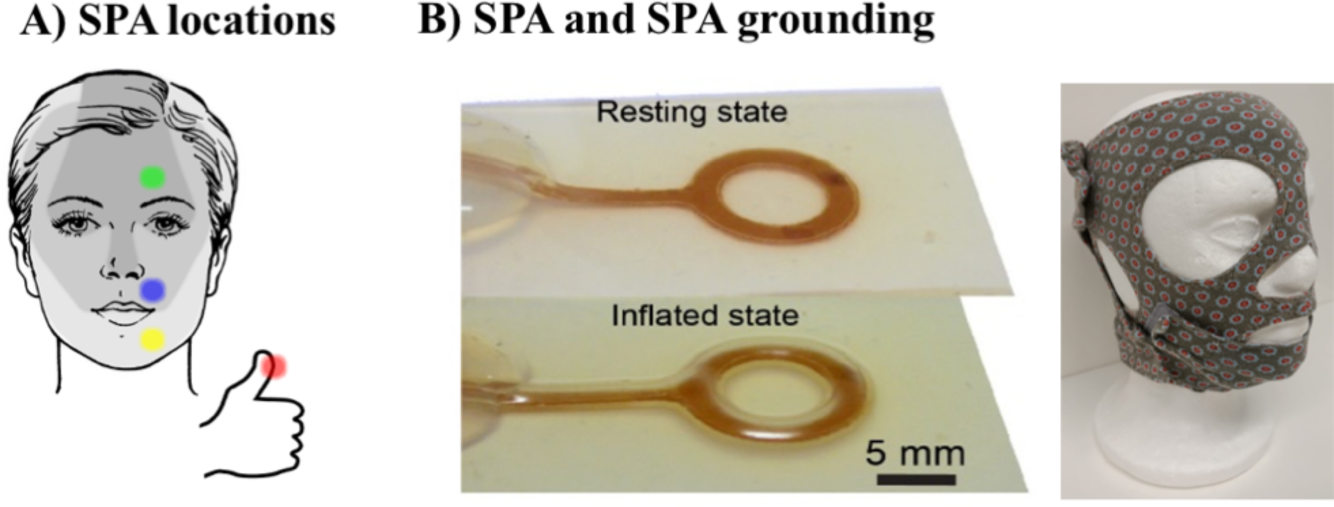
SPA locations and grounding on the face. A) SPAs were attached to the forehead, upper lip, chin, and thumb. We either tested the right or the left side of the face and the right of the left thumb. Shaded areas on the face indicate the skin areas innervated by different branches of the trigeminal nerve. B) SPAs were grounded to the face using a custom made fabric mask.

For any robotic system, a sensor feedback plays an important role to ensure accurate control and understanding of the surrounding environment. The traditionally available pressure or force sensors, needed for providing coherent tactile feedback, work well but like rigid tactile stimulators, these are also limited in their application when they need to be worn by humans, as they often lack the mechanical compliance with the human body. This mismatch obliterates the accurate control of any wearable devices because it is impossible to predict how effective the signal, force, and location of the stimulation is when there is inconsistent and arbitrary grounding. We have hence developed and integrated possibilities of sensing layers in the SPA-skin that provide a localised measurement of interactive forces by virtue of different sensing technologies including an active piezo charge sensing (Sonar and Paik, 2016) or passive methods like resistive strain sensor (Sonar et al., 2019) and soft capacitive skin (Sonar et al., 2018). The sensing layer and the SPA-skin design process tackle this challenge using a finite element based simulation (Agarwal et al., 2017; Moseley et al., 2016) and closed-loop feedback control through the sensing layer. This ensures an accurate tactile stimulation even under variable loading conditions or manufacturing variabilities (Sonar and Paik, 2016).

For the fMRI compliant tactile stimulator application, we designed and prototyped a ring-shaped SPA-skin actuation point (with a 1.4cm diameter and 2mm inlet tube channel) to maximize the application area on the skin while minimizing the volume of air being transferred through the long tubing that helps improve the maximum actuation frequency or bandwidth. This state-of-the-art setup independently controls 8 channels that extended to different SPA- skins. The need for a long tubing (< 6m) limited the volume flow of the air to the actuators during inflation and then back during the deflation cycle. A careful system design was therefore carried out for each component of the pneumatic supply system (PSS) to achieve maximum bandwidth at 6m tube length and maintain the expected functionality of SPAs at less than 80kPa, as discussed in detail in (Joshi et al., 2021; Joshi and Paik, 2021). This allowed us to stretch the actuation bandwidth up to 15Hz in the current setup. When attempting to reach a bandwidth of 30Hz, we were still able to measure 0.5N of force at 30Hz, which is largely above the perception threshold for the face and fingers.

The SPA-skin was pneumatically controlled through a customized control circuit that was interfaced through serial communication to the stimulus computer (see Figure 1). The pneumatic controller was composed of the following components: a stimulus computer, a portable air compressor, a portable pneumatic control circuit module, and corresponding pneumatic tubes. An air compressor (Implotex, Germany) produced the airflow to a pressure regulator (ITV1050, SMC Corp., USA) that was controlled by the microcontroller in the control circuit. The stimulation intensity could be adjusted using a single pressure regulator and an array of high-speed solenoid valves which were driven through high-power MOSFET switches and stimulus from the micro-controller.

A portable pneumatic supply line, AC power supply and USB cable for serial communication enabled the primary extension of the controller setup to be placed just outside the RF shielding walls of the fMRI scanner. A 50mm diameter hole in the wall then carried the 5-6m long and 4mm thick tubes to the SPAs that were places on the face or the fingers. Once connected in this fashion, the system acted as a plug-n-play device that can be easily controlled via the stimulus computer using simple serial commands to turn a given channel on or off at a given frequency. The pressure regulator could be manipulated through the stimulus computer and the working pressure of the pneumatic system could be read in real time. The build-in limitation of 100kPa operational pressure ensures a delamination free operation of the SPA- skin.

### MR interference artefact testing

MR interference artefacts were not expected since all materials inside the MRI scanner room were MR safe. Nevertheless, for the sake of validation, we conducted pilot testing to ensure that no MRI interference artefacts would be induced when the device was turned on with the SPA-skins attached to a water-filled phantom bottle placed inside the MR head coil. To test for artefacts we used the same sequence as used during the fMRI acquisition (see “MRI acquisition” section). As expected, no MR interference artefacts were observed and the presence of the tubing and SPAs inside the scanner room could not be detected in the MRI images.

### Experimental paradigm

The SPA-skins were attached to the forehead (∼1cm above the eyebrow), above the upper lip, and on the chin (see Figure 2A). These sites were chosen to ensure that an SPA-skin was placed on each of the trigeminal nerves’ innervated skin areas. Care was taken to not place SPA-skins on skin areas that are on the border of trigeminal nerve innervations. To ensure good grounding of the SPAs on the face, we placed in-house 3D printed plates on top of the SPA-skins and used a custom-made fabric face mask to apply light pressure to the SPA-skins placed on the face (see Figure 2B). Five further SPA-skins were attached to the fingertips of the left hand using adhesive tape.

Participants viewed a visual display positioned at the head of the scanner bore through a mirror mounted on the head coil. Participants 1-9 were presented with horizontally (for the finger stimulation runs) or vertically (for the face stimulation runs) aligned white circles, corresponding to the different stimulation locations. To cue the participant which location would be stimulated, the circle corresponding to this location turned red 0.8s prior to stimulation onset and remained red until stimulation offset. Participants 10-17 were cued with the words “Forehead”, “Lips”, “Chin”, and “Thumb” in white centred on a black screen. To cue the participant which location would be stimulated, the text cue appeared on the screen 0.8s prior to stimulation onset and remained on the screen until stimulation offset.

Participants were instructed to attend to the highlighted stimulation location as long as the cue was on the screen. Stimulation was presented for 8s at 8Hz with 400ms bursts of stimulation ‘on’ periods followed by a 100ms ‘off’ period to minimise peripheral adaptation. To ensure stable attention during the fMRI runs, stimulations were interrupted in a small percentage of the stimulation blocks (10-20% of trails per run). In these interrupted stimulation trials, stimulation was provided for 4s, after which a 2s silent period was introduced, following by another 2s of stimulation. Care was taken to ensure that the interrupted stimulation trials were equally distributed across the stimulation locations within each run. Participants were instructed to count the number of interrupted stimulation trials and verbally report this at the end of each run.

Since a stronger sensation is expected to lead to a stronger BOLD response, we aimed to match the sensation intensity across stimulation locations prior to the fMRI runs. A stimulation intensity matching task was carried out for the face and finger stimulation runs separately. First, participants were asked to set the optimal stimulation intensity for a reference location. The forehead was chosen as the reference location for the face stimulation runs, and the little finger was chosen as the reference location for the finger stimulation runs. We chose these stimulation locations as references since they are innervated by least mechanoreceptors and can be assumed to be least sensitive (Corniani and Saal, 2020). Participants were instructed that an optimal stimulation intensity would be as strong as possible while remaining focal (i.e., no spread to skin locations not directly underneath the stimulator), comfortable, and stable over the 8s stimulation period (i.e., minimal peripheral adaptation). Participants were asked to respond by means of a button box whether the intensity should be decreased, increased, or should not change. Based on the participants’ responses the air pressure provided to the SPAs was decreased or increased in steps of 5 kPa (leading to a lower or higher stimulation intensity, respectively) or remained stable. If the participant responded twice in a row that the stimulation intensity should not be changed, then the stimulation pressure for the reference location was set at this level for all fMRI tasks. Since the SPAs become fragile when using an air pressure exceeding 80 kPa, this was the maximum pressure setting that could be chosen by participants.

The minimum air pressure provided was set at 20 kPa above atmospheric pressure. Once the optimal stimulation intensity was chosen for the reference location, participants were asked to match the stimulation intensity for the other stimulation locations to the stimulation intensity of the reference location. To enable this matching, participants were initially given 8s of stimulation on the reference location, immediately followed by stimulation of one of the other stimulation locations. Participants were instructed to change the stimulation intensity of the 2^nd^ location to match the stimulation intensity of the reference location as closely as possible. For the face stimulation runs, participants were instructed to match the stimulation intensity of the lips, chin, and thumb to the reference forehead stimulation intensity. For the face stimulation runs, participants were instructed to match the stimulation intensity of the thumb, index, middle, and ring finger to the reference little finger stimulation intensity. As before, if the participant responded twice that the stimulation intensity should not be changed, the stimulation pressure for this stimulation location was set at this level for all fMRI tasks.

Instructions and stimulations were delivered using Psychtoolbox (v3) implemented in Matlab (v2014b). Matlab then communicated with the Arduino board implemented in the SPA controller set-up via a serial port over a proprietary protocol. Head motion was minimized using over-ear MRI-safe headphones or padded cushions.

### Fingers stimulation runs

The finger stimulation blocked design consisted of six conditions: Stimulation conditions for each of the five fingers of the left hand and a rest (no stimulation) condition. The visual cue during the finger stimulation blocks was as described above and the presentation of the word “Rest” indicated the rest condition. Each of these six conditions had a block duration of 8.8s and was repeated five times per run in a counterbalanced order. Each run comprised a different block order and had a duration of 4min and 39.4s. We acquired four blocked design runs, with a total duration of 18min and 37.6s.

### Face stimulation runs

All face stimulation runs involved stimulation of the forehead, lips, chin, and thumb. 14 participants were stimulated on the left side of their face and the left thumb. Three participants were stimulated on the right side of the face and their right thumb. We included thumb stimulation in these runs as the thumb representation borders the face area in S1 (Kikkert et al., 2021, 2016; Kolasinski et al., 2016; Penfield and Rasmussen, 1950; Roux et al., 2018). To uncover face somatotopy we used a blocked design consisting of five conditions: Stimulation conditions for each of the three face locations and the thumb, as well as a rest (no stimulation) condition. The visual cue during the face stimulation blocks were as described above and the presentation of the word “Rest” indicated the rest condition. Each of these five conditions had a block duration of 8.8s and was repeated 8 times per run in a counterbalanced order. Each run comprised a different block order and had a duration of 6min and 7.4s. We acquired four blocked design runs, with a total duration of 24min and 29.6s.

### MRI acquisition

MRI data was acquired using a Philips 3 tesla Ingenia system (Best, The Netherlands). Data of participants 1-9 was collected using a 32-channel head coil and data of participants 10-18 was collected using a 15-channel head coil. fMRI data was acquired using an echo-planar-imaging (EPI) sequence with partial brain coverage: 36 sagittal slices were centred on the postcentral gyrus with coverage over the thalamus and brainstem. We used the following parameters: 2.3mm^3^ spatial resolution, TR: 2200ms, TE: 30ms, flip angle: 82°, SENSE factor: 2.1, 36 slices. We acquired 127 and 167 volumes for the finger and face stimulation runs, respectively. Anatomical T1-weighted images for participants 1-9 were acquired using the following acquisition parameters: TR = 7.7ms, TE = 3.6ms, flip angle = 8°, voxel size = 1mm isotropic, transversal slices = 160. Anatomical T1-weighted images for participants 10-17 were acquired using the following acquisition parameters: 0.7mm^3^ spatial resolution, TR: 9.3ms, TE: 4.4ms, flip angle: 8°.

### MRI analysis

fMRI analysis was implemented using tools from FSL v6.0 (https://fsl.fmrib.ox.ac.uk/fsl/fslwiki) in combination with the RSA toolbox (Nili et al., 2014; Wesselink and Maimon-Mor, 2017) and in-house scripts developed using Matlab (R2018a). Cortical surface analysis and visualisations were realised using Freesurfer (https://surfer.nmr.mgh.harvard.edu/; Dale et al., 1999; Fischl et al., 2001).

#### Preprocessing and image coregistration

Common preprocessing steps were applied to each individual fMRI run using FSL’s Expert Analysis Tool FEAT (version 6.0; https://fsl.fmrib.ox.ac.uk/fsl/fslwiki/FEAT). The following preprocessing steps were included: motion correction using MCFLIRT (Jenkinson et al., 2002), brain extraction using automated brain extraction tool BET (Smith, 2002), spatial smoothing using a 2.3mm full-width-at-half-maximum (FWHM) Gaussian kernel, and high-pass temporal filtering using a cut-off of 90s.

Image coregistration was done in separate, visually inspected, steps. For each participant, a midspace (i.e., an average space in which images are minimally reoriented) was calculated between the 4 face stimulation blocked design runs and, if the fingers were also tested, between the four finger stimulation blocked design runs. We then transformed all fMRI data to these functional midspaces using purely rigid probability mapping in ANTs. Next, we registered each participant’s midspace to the T1-weighted image, initially using 7 degrees of freedom and the mutual information cost function, and then optimised using boundary based registration (BBR; Greve and Fischl, 2009). Each coregistration step was visually inspected and, if needed, manually optimised using blink comparison in Freeview. Structural images were transformed to Montreal Neurological Institute (MNI) standard space using nonlinear registration (FNIRT).

#### Univariate analysis

First-level parameter estimates were computed using a voxel-based general linear model (GLM) based on the gamma hemodynamic response function and its temporal derivatives. Time series statistical analysis was carried out using FILM (FMRIB’s Improved Linear Model) with local autocorrelation correction. To reduce noise artefacts, CSF and WM scan wise time series were added to the model as nuisance regressors. Data were further assessed for excessive motion, and volumes with an estimated absolute mean displacement > 1.15mm (half of the functional voxel size) were scrubbed. Contrasts were defined for each stimulation condition versus rest, for overall face or finger stimulation conditions versus rest, and for each stimulation condition versus all other stimulation conditions. We then used a fixed-effects higher-level analysis to average across the finger stimulation runs and across the face stimulation runs separately for each individual participant. Z-statistic images were thresholded using clusters determined by Z > 2.3 and p < .05 family-wise-error-corrected cluster significance thresholding was applied.

To visualise inter-participant consistency of somatotopic finger and face selective representations, we calculated cortical activation probability maps. Cortical surface projections were constructed from participants’ T1-weighted images. Each participant’s cortical surface was inflated into a sphere and aligned to the Freesurfer 2D average atlas using sulcal depth and curvature information. The thresholded stimulation site selective (i.e., condition versus all other stimulation conditions) contrast maps from each participant’s fixed effects higher-level analysis were resampled to the Freesurfer 2D average atlas and binarized. We then calculated finger-specific and face part-specific inter-participant probability maps.

Lastly, to visualise which brain areas were activated during overall face and fingers stimulation, whole-brain group averages were assessed for the stimulation of all fingers versus rest and stimulation of all face locations versus rest contrasts. Group-level analysis was performed for the finger and face stimulation runs separately using FMRIB’s Local Analysis of Mixed Effects (Woolrich et al., 2004). Data collected for individuals of whom we tested the right side of the face was flipped on the midsagittal plane before conducting group analysis to ensure that the tested hemisphere was consistently aligned. Z-statistic images were thresholded using clusters determined by Z > 3.1 and p < .05 family-wise-error-corrected cluster significance thresholding was applied.

#### Representational similarity analysis

A fuller description of somatotopic representations can be obtained by taking into account the entire fine-grained activity pattern of stimulated fingers or face parts. Representational dissimilarity was therefore estimated between fingers or between face locations using the cross-validated squared Mahalanobis distance (or crossnobis distance; Nili et al., 2014; Wesselink and Maimon-Mor, 2017b). We closely followed previously described procedures (Kikkert et al., 2021; Wesselink et al., 2019).

We first defined an anatomical S1 hand and an S1 face ROI using the probabilistic Brodmann area parcellation provided by recon-all in Freesurfer. Each participant’s cortical parcellations for Brodmann areas 1, 2, 3a and 3b were converted to volumetric space and merged to form a single S1 mask. Next, any holes in these converted masks were filled and the non-zero voxels in the mask were mean dilated. We restricted this S1 mask such that it spanned a 2cm strip medial/lateral to the anatomical location of the hand knob to form the S1 hand ROI (Yousry et al., 1997). The S1 area inferior to this S1 hand ROI was then extracted to form the S1 face ROI.

We computed the dissimilarity between the activity patterns measured for each pair of stimulation conditions within the S1 hand or S1 face ROI for the fingers and face stimulation runs, respectively. We extracted the voxel-wise parameter estimates (betas) and the model fit residuals under the ROI and prewhitened the betas using the model fit residuals. We then calculated the cross-validated squared Mahalanobis distances between each pair of conditions, using the four runs as independent cross-validation folds, and averaged the resulting distances across the folds.

The dissimilarity values for all pairs of conditions were initially assembled in a representational dissimilarity matrix (RDM), with a width and height corresponding to the number of conditions (i.e., a 5x5 RDM for the finger stimulation runs and a 4x4 RDM for the face stimulation runs). Since the RDM is mirrored across the diagonal with meaningless 0’s on the diagonal, all statistical analysis was conducted on the unique values of the RDM (10 unique values for the finger stimulation runs and six unique values for the face stimulation runs). Finally, we performed multidimensional scaling (MDS) to visualise the dissimilarity structure of the RDM in an intuitive manner. MDS projects the higher-dimensional RDM into a lower-dimensional space while preserving the inter-condition dissimilarity values as well as possible (Borg and Groenen, 2005). MDS was performed for each individual participant and then averaged per group after Procrustes alignment to remove arbitrary rotation induced by MDS.

We estimated the strength of the representation or “representational separability” by averaging the unique off-diagonal values of the RDM. If it is impossible to statistically differentiate between conditions (i.e., when a parameter is not represented in the ROI), the expected value of the distance estimate would be 0. If it is possible to distinguish between activity patterns this value will be larger than 0. To further ensure that our S1 face ROIs contained face information, we created a cerebral spinal fluid (CSF) ROI that would not contain somatotopic face information. We then repeated our face RSA analysis in this ROI and statistically compared the separability of the CSF ROI to the separability in the S1 face ROI.

### Statistics

Statistical analysis was carried out using SPSS (v25). Non-parametric statistical testing was used for analysis of the finger stimulation data. Parametric statistical analysis was carried out for the face stimulation data after checking for normality using the Shapiro-wilk test. Non-parametric statistical testing in case of normality violations. All testing was two-tailed and we used the Benjamini-Hochberg procedure to control the false discovery rate with q < 0.05.

### Data availability

Full details of the experimental protocol are available on clinicaltrials.gov under the number NCT03772548. Data is shared on <link will be made available upon publication>.

## Results

In the current study, we aimed to validate the use of our newly developed SPA-setup for somatotopic mapping using fMRI. We first explored whether SPA tactile stimulation would elicit characteristic finger representations in S1. Next, we used our SPA setup to uncover the somatotopic layout of the face in S1.

### Vibrotactile SPA-skin stimulation induces activity in somatosensory processing areas

We first examined task-related brain activity elicited by tactile stimulation applied to the fingers (Figure 3A) and the face (Figure 3B) using our SPA-skin setup. As expected, we found that both stimulation on the fingers and the face activated S1 and secondary somatosensory cortex (S2) contralateral to the stimulation site. Ipsilateral S1 and S2 activity was less widespread for both the fingers and face stimulation. Note that a direct comparison with regards to the level of activity between the finger and face stimulation runs is challenging given that activity elicited by fingers stimulation was only tested on a subset of eight participants and seventeen participants were tested to examine activity elicited by face stimulation.

**Figure 3:**
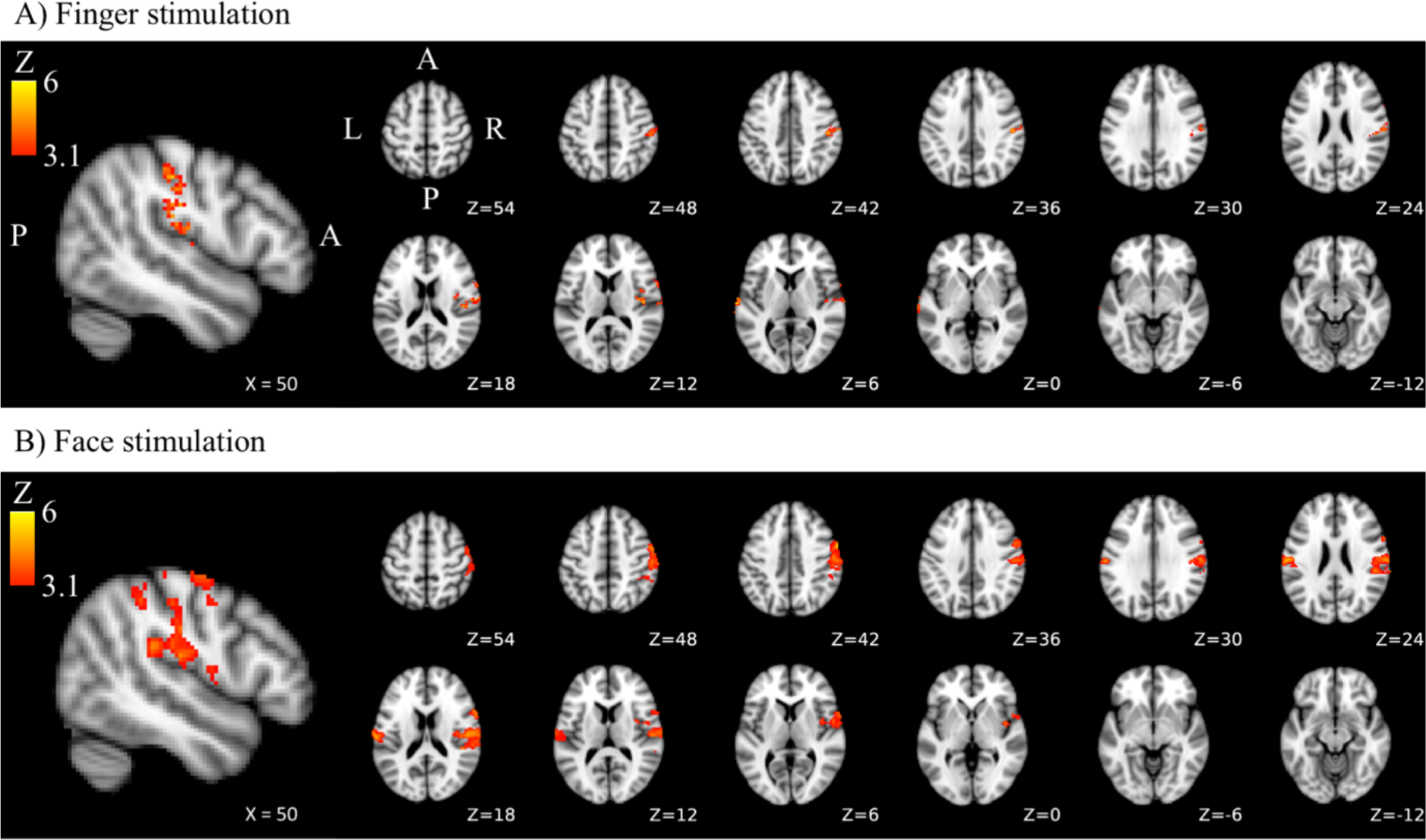
Brain activity in the contralateral hemisphere during sensory stimulation using SPA technology. Both fingers (A) and face (B) stimulation elicited activity in S1 and secondary somatosensory cortex. L = left; R = right; P = posterior, A = anterior.

### Finger representations

We first explored finger selective representations. We contrasted activity elicited during each finger stimulation against activity during stimulation of all other fingers and binarized the resulting finger selective maps. We then created inter-participant probability maps representing the number of participants exhibiting specific finger selectivity (i.e., using a winner-take-all approach) for each vertex in S1. These maps exhibited, as expected (Kolasinski et al., 2016), that finger selectivity was not perfectly consistent across participants (Figure 4A). We did however observe a characteristic gradient of finger preference progressing from the thumb, for which inter-subject probability was highest lateral of the S1 hand area, to the little finger, for which inter-subject probability was highest medial of the S1 hand area. Qualitative inspection suggests that inter-participant consistency was highest for the thumb and lowest for the little finger representation.

**Figure 4:**
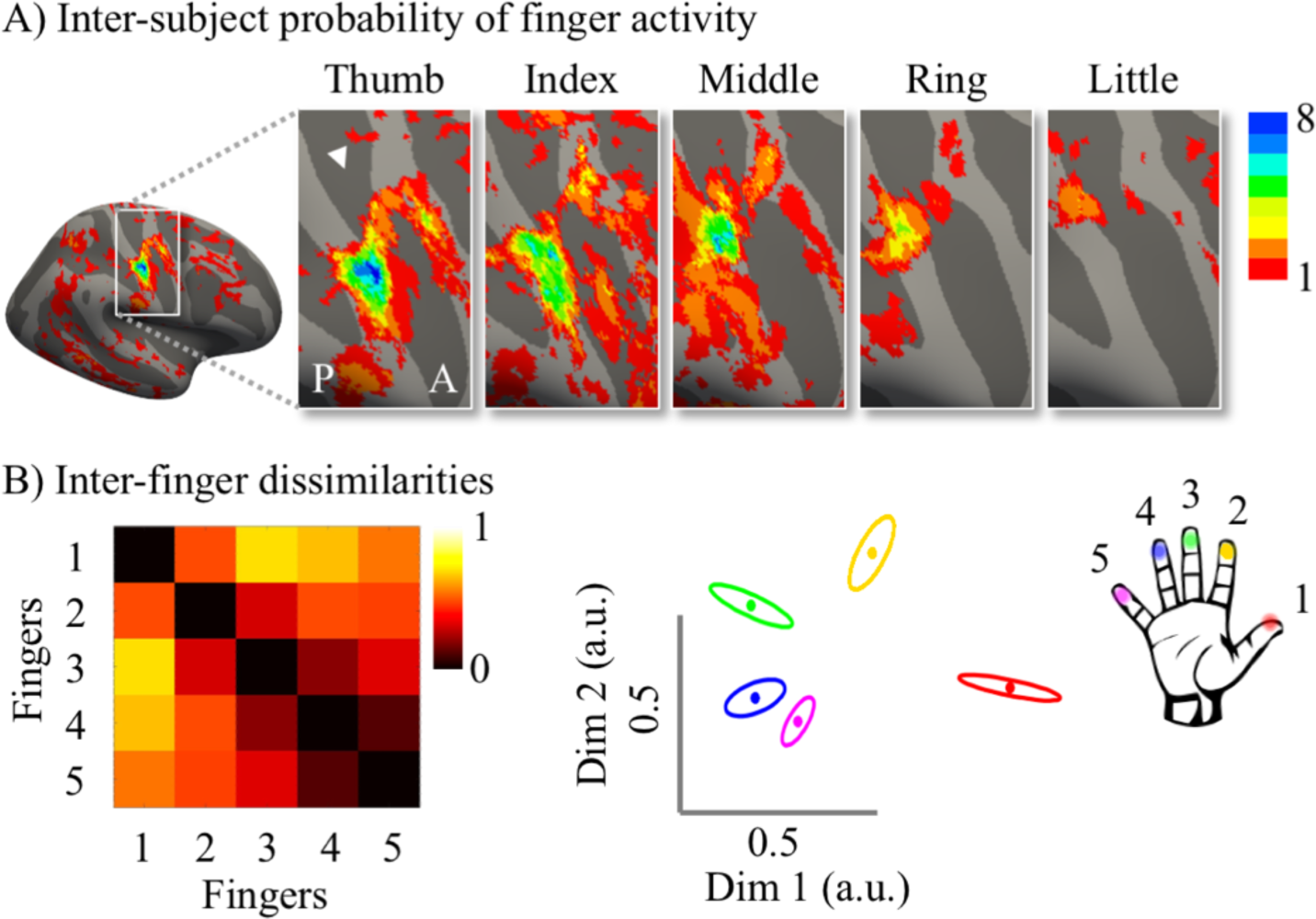
SPA stimulation uncovers typical finger somatotopy. A) inter-participant probability maps of finger selective representations. Colours indicate the number of participants (ranging from 1 (red) to 8 (blue)) who demonstrated finger selectivity for a given vertex. Typical finger selectivity is characterised by a progression of finger selectivity from the thumb (laterally) to the little finger (medially). We observed a characteristic gradient of finger preference progressing from the thumb, for which inter-subject probability was highest lateral of the S1 hand area, to the little finger, for which inter-subject probability was highest medial of the S1 hand area. Qualitative inspection suggests that inter-participant consistency was lowest for the little finger representation and highest for the thumb representation. The white arrow indicates the central sulcus. A = anterior; P = posterior. B) The representational structure of inter-finger distances. Left: Representational Dissimilarity Matrix (RDM). Individual stimulation sites are represented by numbers: thumb = 1; index finger = 2; middle finger = 3; ring finger = 4; little finger = 5. Right: 2-dimensional projection of the RDM. Inter-finger distance is reflected by the distance in the two dimensions. Individual fingers are represented by different colours: thumb = red; index finger = yellow; middle finger = green; ring finger = blue; little finger = purple. By stimulating fingers using SPA technology, we found a classical and frequently reported inter-finger distance pattern in the hand area of S1. Ellipses represent the between-participants standard error after Procrustes alignment.

Next, we explored a 2^nd^ somatotopic principle that is known to be consistent across individuals: the pattern of distances between finger representations. We examined this intricate relationship between finger representations using representational dissimilarity analysis. We found a typical pattern of inter-finger representational distances where neighbouring fingers have relatively lower representational distances compared to fingers that are further apart. Furthermore, fingers that we use more frequently together in daily life (e.g., the ring and little fingers) had lower representational distances compared to fingers we use more separately in daily life (e.g., the thumb and index finger (Ejaz et al., 2015)). This inter-finger representational distance pattern found using SPA tactile stimulation of fingers (Figure 4B) was highly similar to the inter-finger representational distance patterns that we and others have described previously in fMRI experiments using individual finger movements (Ejaz et al., 2015; Kikkert et al., 2021, 2016; Sanders et al., 2019; Wesselink et al., 2019) or individual finger tactile stimulation (Sanders et al., 2019). The inter-finger representational distances were averaged across finger pairs within each participant to obtain an estimate for average inter-finger representational separability (see Figure 5C), or ‘representation strength’. We found that inter-finger separability in the S1 hand area was greater than 0 (Z = -2.52, p = 0.01), showing that the S1 hand area contained information about individual finger representations. Furthermore, there was significant greater separability in the S1 hand area compared to a control cerebral spinal fluid (CSF) ROI that would not be expected to contain finger specific information (Z = -2.52, p = 0.01). Together, our results demonstrate that our SPA setup can be used to reliably map the well characterised somatotopic layout of the hand.

**Figure 5:**
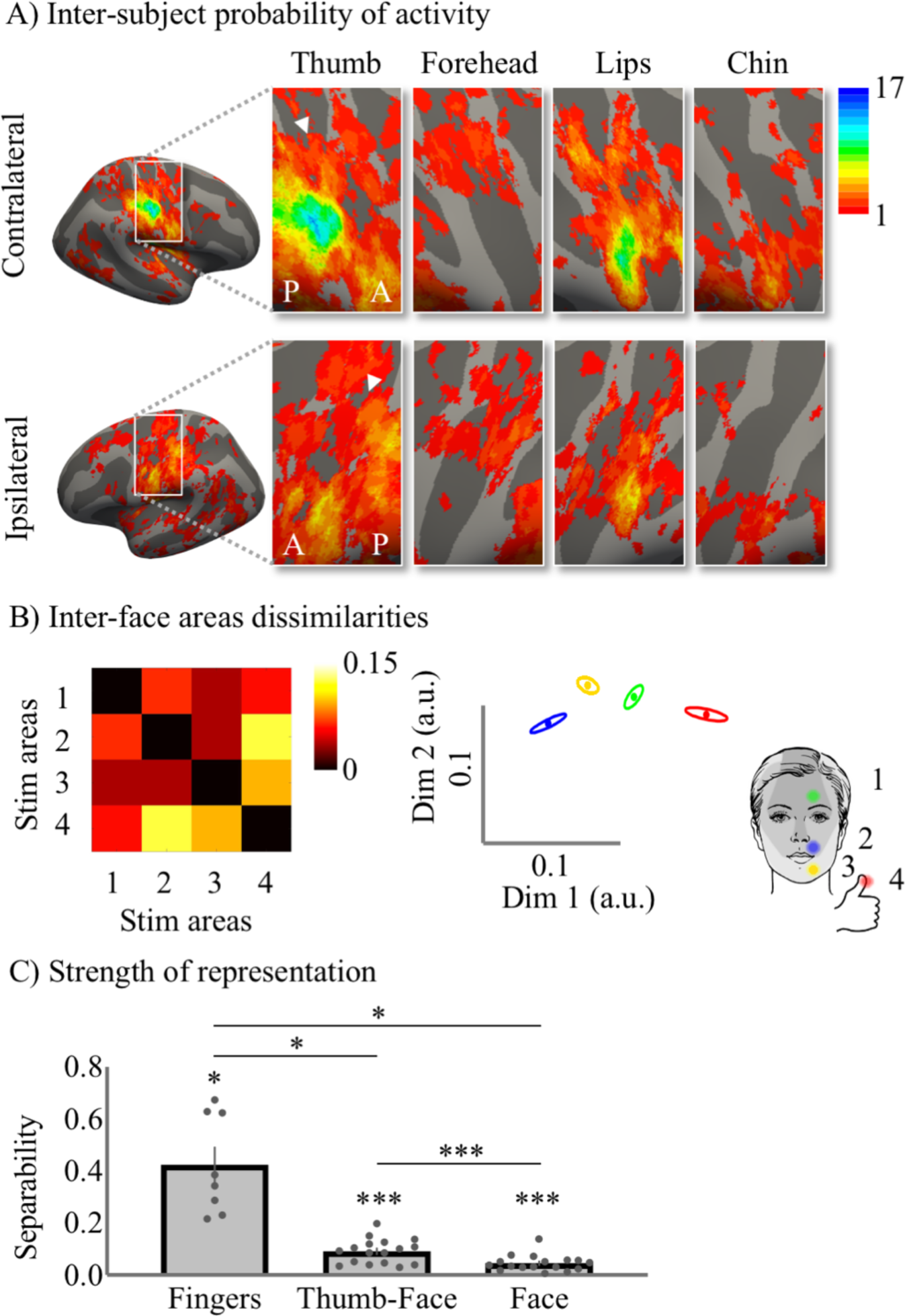
Face somatotopy uncovered using SPA stimulation. A) Inter-participant somatotopic probability face maps. Colours indicate the number of participants (ranging from 1 (red) to 17 (blue)) who demonstrated face part or thumb selectivity for a given vertex (i.e., stronger activity compared to the other stimulation sites). Qualitative inspection suggests that inter-participant consistency contralaterally was lowest for the forehead representation and highest for the thumb representation. Furthermore, inter-participant consistency was higher in contralateral, compared to ipsilateral S1. White arrows indicate the central sulcus. A = anterior; P = posterior. B) The representational structure of inter-stimulation site distances. Left: Representational Dissimilarity Matrix (RDM). Individual stimulation sites are represented by numbers: forehead = 1; upper lip = 2; chin = 3; thumb = 4. Right: 2-dimensional projection of the RDM. Inter-stimulation site distance is reflected by the distance in the two dimensions. Individual stimulation sites are represented by different colours: forehead = green; upper lip = blue; chin = yellow; thumb = red. Ellipses represent the between-participants standard error after Procrustes alignment. C) Separability (i.e., average dissimilarity), or ‘representation strength’, for fingers-to-fingers in the S1 hand area, and thumb-to-face and face-to-face in the S1 face area. There was significant separability (i.e., greater than 0) for fingers-to-fingers in the S1 hand area, as well as for thumb-to-face and face-to-face in the S1 face area. Inter-fingers separability in the S1 hand area was greater than thumb-to-face and face-to-face separability in the S1 face area. Lastly, thumb-to-face separability in the S1 face area was higher than face-to-face separability in the S1 face area. * = p < 0.05; *** = p < 0.001.

### Face representations

One advantage of our SPA setup is that, unlike most commercially available vibrotactile devices, it is able to provide vibrotactile stimulation to localised areas of the face (i.e., in the MRI head coil) without causing any imaging artefacts or safety issues. We made use of this advantage by examining the somatotopic layout of the face using our SPA setup. First, as for the fingers, we created inter-participant probability maps of forehead, lip, chin, and thumb selective representations (Figure 5A). The resulting inter-participant probability maps represent the number of participants exhibiting selectivity (i.e., using a winner-take-all principle) for a specific stimulation site for each vertex in S1. Qualitative inspection suggests that the selective thumb and lip activity was more consistent across participants compared to the forehead and chin. Furthermore, stimulation site selective activity was more consistent across participants in the contralateral compared to the ipsilateral hemisphere.

We then used RSA in the S1 face area to investigate representational distances between stimulation sites (i.e., between the forehead, lip, chin and thumb) that were stimulated during the face stimulation runs (Figure 5B). The averaged representational distances (i.e., separability) between face sites (i.e., excluding the face-thumb distances) were markedly lower compared to the representational distances found between fingers in the S1 hand area (Figure 5C; Z = -2.52, p = 0.01) and between the thumb and face sites in the S1 face area (Figure 5C; t(16) = 6.11, p < 0.001). However, we found that face-face site separability in the S1 face area was greater than 0 (Figure 5C; t(16) = 6.10, p < 0.001) and greater than separability in a control CSF ROI (t(16) = -6.00, p < 0.001), indicating that the S1 face area contained information about face sites. We found a clear pattern of inter-face site representational distances in the S1 face area revealing that the forehead had the lowest representational distance from the thumb. The chin had more representational distance from the thumb than the forehead, and the lips were most distant from the thumb.

## Discussion

Providing somatosensory stimulation in an MR environment is challenging. Our results show that somatosensory representations can be mapped using new SPA technology that is MR-compatible and can be used within the MR head coil without causing artefacts. Using our SPA setup, we found a typical somatotopic layout of participants’ fingers, both in terms of the gradient of finger selectivity and the inter-finger representational distance pattern. This validates the use of our SPA setup for mapping somatotopic representations.

We further probed face somatotopy using automated tactile stimulation to the forehead, lips, and chin – 3 face sites that are innervated by different branches of the trigeminal nerve. Our univariate analysis revealed a great degree of inter-participant variability of face-site selective representations. While the location of the thumb and lip representations were relatively stable across participants, the forehead and chin representations were much less stable. Potentially, the low density of mechanoreceptors in the chin and forehead may have caused their activations to be ‘overshadowed’ by overlapping thumb and lip representations in this winner-take-all analysis. While we attempted to control for any variability in sensed intensity using our intensity matching task, it is likely that the forehead and chin are represented less prominently in S1 compared to the thumb and lip and may therefore not ‘win’ in a winner-take-all univariate stimulation site-selective analysis. Indeed, we suggest that it is more appropriate to study face somatotopy using an analysis method that does not adhere to such a winner-take-all principle and instead is sensitive to representational overlap or representational distances between face parts. Using RSA, we revealed that the forehead was least dissimilar to the thumb representation, followed by the chin representation, and lastly by the lip representation. These results are in line with and extend the results of a recent study using RSA and movements of the forehead, nose, lips, and tongue to study face somatotopy in S1. As in our study, Root et al. (2021) reported that the forehead was least distant from the hand representation. They further found that the nose was representationally least distant from the forehead representation and that the lips were furthest from the forehead representation. Root et al. (2021) could not disentangle the lips and chin representation, due to the nature of their movement task. As such, we extend their results by showing that, in representational space, the chin is in-between the lip and forehead in S1.

The aim of this study was to validate the use of a simple pneumatic vibrotactile device for somatotopic mapping inside an MRI scanner and specifically the head coil. Soft robotics emerged as a new field in robotics over a past decade and has been growing across the applications in medical technologies, personal care, environmental monitoring and entertainment (Bauer et al., 2014; Gibson, 2018; Hawkes et al., 2017; Jafari et al., 2016; Polygerinos et al., 2017). Soft robotics tackles the interactivity and wearability challenges posed to traditional rigid body robotic systems by using soft material based sensors and actuators to replace the traditional sensors and motors. These soft materials enable an inherently safe and mechanically compliant feedback to the environment including a human wearer (Polygerinos et al., 2017, 2015; Pons, 2008; Robertson and Paik, 2017). Wearability of soft technology can further be used to improve traditional haptic communication devices that currently only provide limited wearability due to their bulky and rigid-form factors (Cholewiak and Collins, 2003; Jones, 2011; Yun et al., 2017). Present haptic feedback devices mainly rely on motors and components driven using electromagnetic eccentric mass or piezoelectric motors that have limited wearability due to their bulky rigid mechanisms and cannot be used in environments prone to electromagnetic interference like the MRI environment (Alahakone and Senanayake, 2009; Cholewiak and Collins, 2003; “Smart Tactor Development Kit,” 2021).

While pneumatic tactile stimulators are highly applicable in studies investigating somatotopy, especially when investigating body part representations in or nearby the MR head coil, they generally have a smaller frequency range and more limited actuation control compared to piezoelectric devices. Due to the length of our plastic tubes, our SPA setup can provide accurate control of frequencies up to 35Hz. While this stimulation frequency is reasonable for somatotopic mapping experiments, it may not be ideal when investigating neural responses to stimulation of specific mechanoreceptors. Furthermore, while our SPA-skin system can stimulate suprathreshold, its stimulation amplitude is limited due to the used materials. As such, for studies requiring highly intense sensations on skin surfaces (< 1N) with very low innervation of mechanoreceptors (e.g., on the neck or ankle), another material combination needs to be examined. Furthermore, the pneumatic supply system components (tubing resistance, compressibility and inertia of the oscillating air, conductance of various components) and the extent of grounding force with which the SPA is fixed to the stimulation site and could influence the exact amount of skin indentation.

Despite these limitations of the current system, we could reveal clear somatotopic representational patterns of different body parts by providing vibrotactile stimulation using the SPA-skin technology. Given that in our SPA-skin system all materials in the MRI scanner room are non-metallic, no MR safety and image quality certifications are required when using this device. SPAs are small and can easily be attached to different sites of the body. The SPA setup is flexible, easy-to-implement, precise, portable, fast and offers a cost-effective solution in comparison to commercially available devices that often induce artefacts, are highly expensive, require active shielding, or hardware modifications.

## Author contributions

Conceptualization; Data curation; Formal analysis; Funding acquisition; Investigation; Methodology; Project administration; Resources; Software; Supervision; Validation; Visualization; Roles/Writing - original draft; Writing - review & editing

## Acknowledgements

We thank Lena Salzmann, Lydia Kampf, Nicolin Gauler, Philipp Barzyk, Silvia Hofer, Daniel Reichmuth, and Sijamini Baskaralingam for assistance with piloting or data collection. We thank Daniel Woolley and Charles Lambelet for technical support.

